# A quantised cyclin-based cell cycle model

**DOI:** 10.1101/2020.07.13.200303

**Authors:** Chris Emerson, Lindsey Bennie, Dermot Green, Fred Currell, Jonathan A. Coulter

**Affiliations:** School of Pharmacy, Queens university Belfast, Belfast, Northern Ireland, United Kingdom; School of Mathematics and Physics, Queens University Belfast. Belfast, Northern Ireland, United Kingdom; Department of Chemistry and The Dalton Cumbrian Facility, University of Manchester, Manchester, United Kingdom

## Abstract

Computational modelling is an important research tool, helping predict the outcome of proposed treatment plans or to illuminate the mechanics of tumour growth. *In silico* modelling has been used in every aspect of cancer research from DNA damage and repair, tumour growth, drug/tumour interactions, and mutational status. Indeed, modelling even holds potential in understanding the interactions between individual proteins on a single cell basis. Here, we present a computational model of the cell cycle network of the cyclin family of proteins (cyclin A, B, D and E). This model has been quantised using western blot and flow cytometry data from a synchronised HUVEC line to enable the determination of the absolute number of cyclin protein molecules per cell. This quantification allows the model to have stringent controls over the thresholds between transitions. The results show that the peak values obtained for the four cyclins are similar with cyclin B having a peak values of 5×10^6^ to 9×10^6^ molecules per cell. Comparing this value with the number of actin proteins, 5E^8^, shows that despite their importance, the level of cyclin family proteins are approximately 2 orders of magnitude lower. The efficiency of the model presented would also allow for its use as an internal component in more complex models such as a tumour growth model, in which each individual cell would have its own cell cycle calculated independently from neighbouring cells. Additionally, the model can also be used to help understand the impact of novel therapeutic interventions on cell cycle progression.

**Author Summary:** Protein and gene networks control every physiological behaviour of cells, with the cell cycle being controlled by the network of genes that promote the cyclin family of proteins. These networks hold the key to creating accurate and relevant biological models. Normally these models are presented with relative protein concentrations without any real world counterpart to their outputs. The model presented within shows and advancement of this approach by calculating the absolute concentration of each cyclin protein in one cell as it progresses through the cell cycle. This model employs Boolean variables to represent the genetic network, either the gene is active or not, and continuous variables to represent the concentrations of the proteins. This hybridised approach allows for rapid calculations of the protein concentrations and of the cell cycle progression allowing for a model that could be easily incorporated into larger tumour models, allowing for the tracking of discrete cells within the tumour.

## Introduction

The progress of a eukaryotic cell through the cell cycle is controlled by a complex interaction network of two main protein families, the cyclins and the cyclin-dependent-kinases, CDK’s.^[1–3]^ For regulation of the cell cycle there are four main cyclins and CDK’s; Cyclins A, B, D and E, and CDK’s 1, 2, 4 and 6. Each cyclin is responsible for regulation of different cell cycle phases when complexed with the corresponding CDK. This full interaction network is illustrated in Figure.1 along with the Boolean network to represent the cell cycle.

**Figure. 1.**
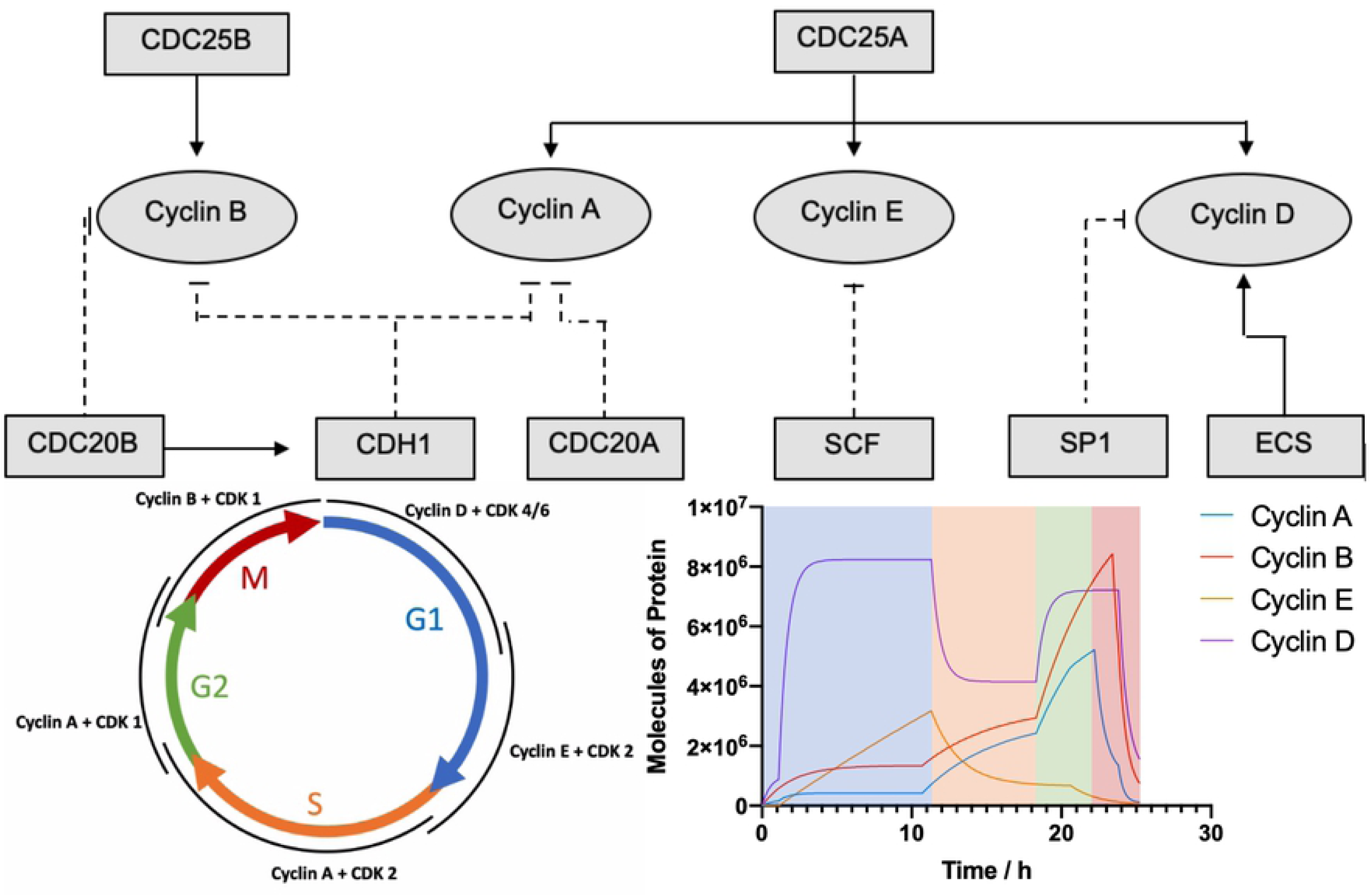
showing the activation pairing of the cyclins and the CDK’s responsible for each of the cell cycle phases(G_1_, S, G_2_ and M) and phase transitions. Alongside the computational output of the model created in this work, colour coded to each phase of the and the Boolean network.

Many computational models of the cell cycle have been created using different mathematical approaches depending on the network or interactions being studied; continuous modelling, Boolean modelling or a combination of both, denoted as hybrid modelling.^[5–7]^ Continuous methods have been used to study the concentrations of proteins across the whole cell cycle, while the Boolean method solves at discrete steps, were real-world time relating to each step is not consistent.^[8]^ Boolean modelling is mainly used to model a network where every variable has only two states either active (defined as 1), or inactive (defined as 0). This on/off state can be used to model a simplistic view of gene expression and therefore expanded to protein translation. However, this approach discards the high level of temporal detail achieved by continuous methods. A powerful example of this is type of modelling is work carried out by Lee *et al.* (2004).^[6]^ Using a Boolean method, the authors discovered that by creating a network of protein interactions, the system would follow a controlled cycle of thirteen unique Boolean states, corresponding to the cell cycle. Boolean methods provide very accurate interaction networks and are often used to rapidly identify stable states. Conversely, continuous methods provide a direct result between the relative levels of variables being studied.^[9]^

Continuous modelling’s main benefit stems from the ability of the model to be solved over any desired time point or series. Therefore, it has repeatedly been used to study variation of concentration gradients within biological systems, including protein concentration, oxygen diffusion and tumour growth.^[10–15]^ An early study investigating protein concentration variation was reported by Tyson *et al.* (1991).^[15]^ The authors developed a model based on the interaction and phosphorylation status of the proteins cdc2 and cyclin in yeast cells. The model investigated the formation of maturation promoting factor (MPF), a protein complex that triggers the entry of yeast cells into mitosis. It uses a single protein complex to model the cell cycle, requiring the solving of six simultaneous partial differential equations. As such the main limitation of a pure continuous model is the number of variables and the computational time required to solve the model.

Hybrid modelling is a combination of both Boolean and continuous models.^[10]^ Hybrid models combine the framework and steps of a Boolean model, however, they also contain unique differential equations which require resolving before progression to the next Boolean step. This approach was adopted by Singhania *et al*. (2011) who used the stable state Boolean network created by Li (2004) to incorporate differential equations for the three main cyclin proteins controlling cell cycle progression.^[5]^

The model presented in this work is an extension of the hybrid model, presented by Singhania *et al* (2011), that combines both approaches, an overarching Boolean network with continuous equations underneath.^[5]^ It allows for the high data output achievable from a continuous model, as well as benefiting from the computational efficiency that a Boolean model gives. However, the model presented here differs in two important ways. Firstly, the inclusion of cyclin D allows for a stricter control of the initial entry into the cell cycle. Secondly the model takes account of the cyclin molecules as discrete entities rather than continuum quantities. Accounting for them in this way allows the stochastic features of the cell cycle to emerge rather than for them to be added in manually. The use of Boolean networks also allows for the differential equations to be solved analytically for any time during a single Boolean phase. This drastically improves the compute time for the cell cycle allowing for the collection of a high volume of results.

Furthermore, by directly comparing the model to biological data collected from a synchronised human umbilical endothelial cells (HUVEC), we aimed to test model accuracy and improve the quality of outputs by quantising it to outputs as tangible units of protein molecules per cell. Cell population synchronisation is a necessary strategy when studying specific phases of the cell cycle. There are numerous experimental strategies used to synchronise cell populations, these include physical separation using FACS (fluorescence-activated cell sorting), chemical blockades, nutrient depravation and contact inhibition.^[16–19]^ To allow the comparison between biological and computational data, chemical based synchronisation offers the cleanest and an easily reproducible synchronisation. Within this study the drug nocodazole was used as it acts as a rapidly-reversable inhibitor of microtubule polymerization, stalling the cycle at the G_2_/M checkpoint.^[20]^ Nocodazole is a synthetic tubulin-binding agent with antineoplastic activity. Binding to beta-tubulin, nocodazole disrupts the microtubule network required for chromosomal rearrangement during mitosis.^[21]^ As such, cells treated with nocodazole fail to enter mitosis with cells accumulating at the G_2_/M checkpoint.^[22]^

## Results

As this model is based on multiple single cells completing the cell cycle, all biological experiments must be performed on synchronised cell populations. This was achieved using nocodazole, as detailed above. However, to ensure that drug induced effects did not influence data outputs, phenotypic nocodazole characterisation was performed, enabling the quantisation of the computational model. First we will discuss the collection of the biological data, then the method of converting the model into absolute concentration.

### Nocodazole toxicity and G_2_/M checkpoint stalling

The biochemical impact of nocodazole is well understood, blocking spindle formation and impeding mitosis. However, nocodazole confers this effect in a strong concentration dependent manner^22^. Thus, for the HUVEC line differing nocodazole concentrations were investigated for both cytotoxicity and <u>G_2_/M</u>stalling efficiency.^[20]^ Survival following nocodazole treatment was normalised against untreated cells and plotted as a surviving fraction. Figure 2 demonstrates clear dose dependent toxicity of nocodazole up to 200 μg/ml.

**Figure. 2.**
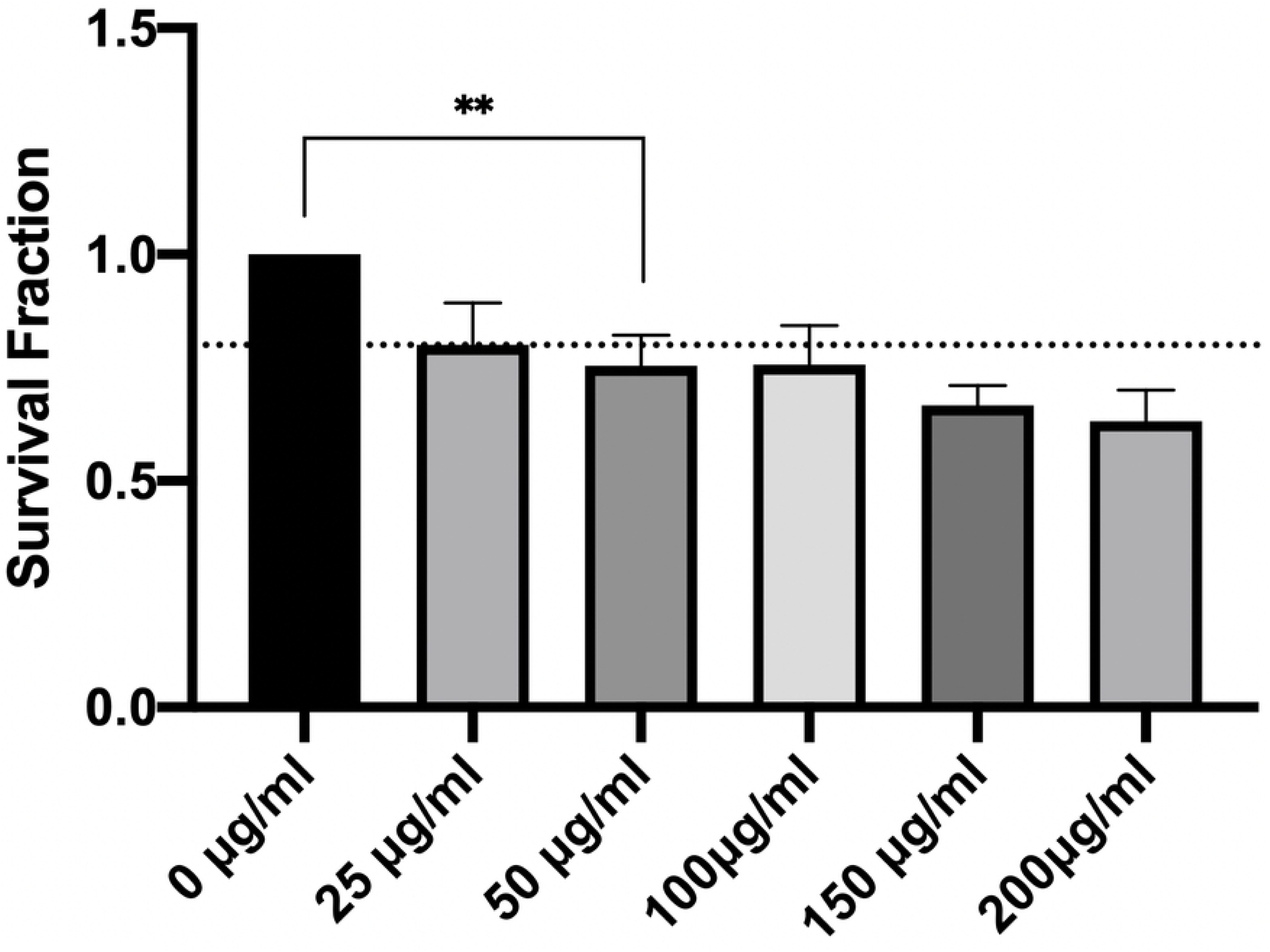
Concentration dependent toxicity of nocodazole in HUVEC’s. Drug concentrations in excess of 50 μg/mL reduced survival by >20%, representing weak direct cytotoxicity as defined by ISO 10993-5. Statistical analysis: one-way ANOVA p=0.0032

The International Organization for Standardization (ISO) standards 10993-5 define multiple categories of cytotoxicity.^[23]^ Direct cell death defined as triggering between 20% - 40% reduction in survival is classified as weakly cytotoxic, from 40% - 60% as moderately cytotoxic and above 60% is strongly cytotoxic. Applying these conventions, nocodazole exhibits weak cytotoxicity at all concentrations tested with survival reduced to 63% relative to untreated controls at the highest concentration (200 μg/ml) tested. Nevertheless to mitigate potential direct toxicity effects, a deeper understanding of the stalling efficiency was studied using flow cytometry (Figure 3). Combining the results from figures 2 & 3 allows for a correlation between the toxic effects and the stalling efficiency at a variety of selected concentrations.

**Figure 3.**
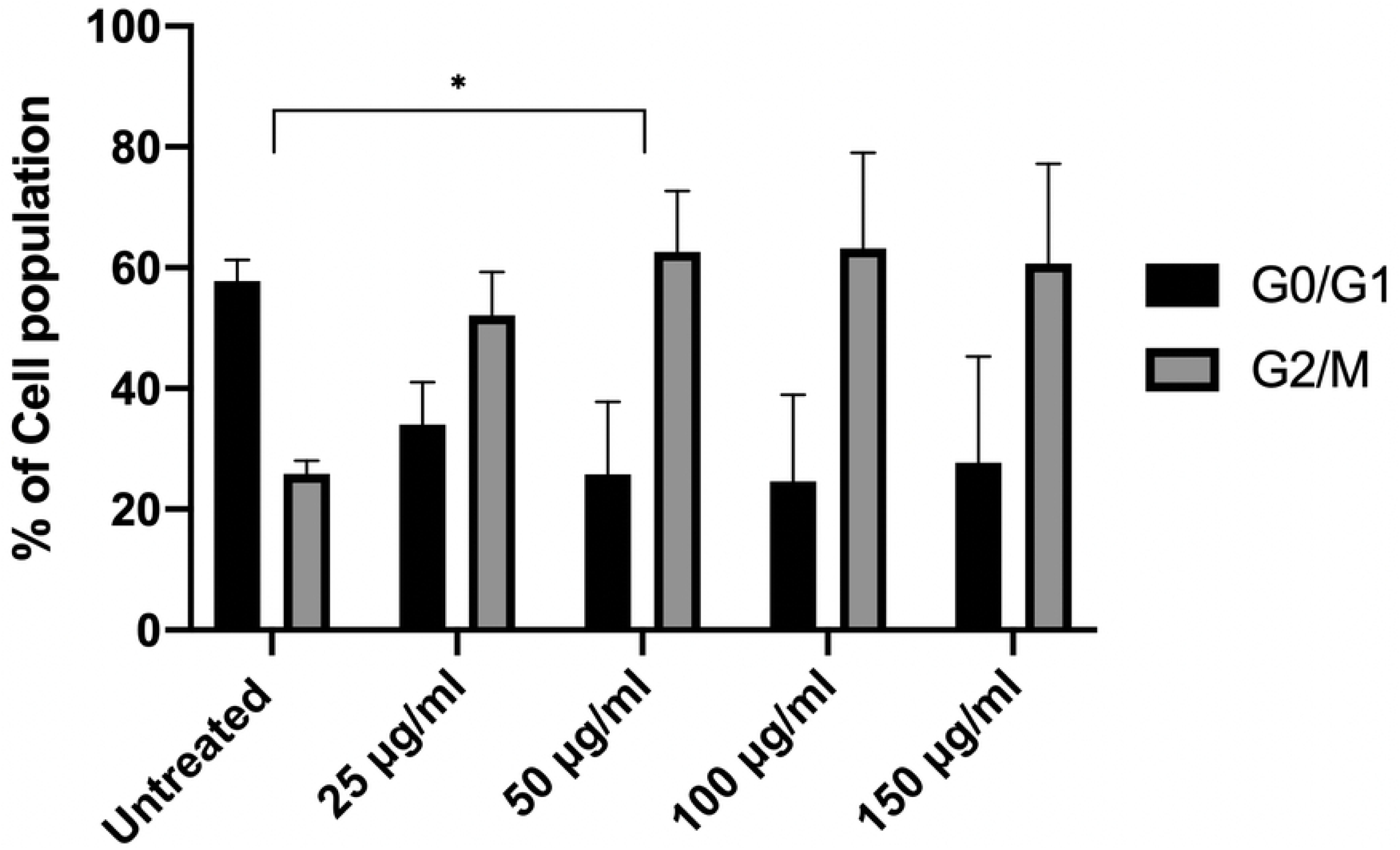
Concentration dependent G_2_/M checkpoint stalling by nocodazole. Maximum cell cycle stalling was achieved using nocodazole at 50 μg/mL, while minimising the toxicity risk. Statistically (one-way ANOVA), nocodazole concentration >50 μg/ml caused significant (p<0.05) G_2_/M arrest.

Nocodazole concentrations in excess of 50 μg/mL confer no additional advantage to increasing the relative fraction of stalled cells. As such 50 μg/mL was selected for all subsequent experiments. To examine the impact of various interventions on cell cycle, it is also essential to understand the dynamics of cell cycle progression following the removal of nocodazole blockade. We therefore preformed flow cytometry via propidium iodide staining, allowing the measurement of the cell cycle dynamics. This data provides the time offset or the t=0 value (time correction factor) when drawing comparisons between the *in vitro* data and the *in silico* data.

Figure 4 shows that the percentage of the cells in G_2_/M is almost equal between 1.5 and 3.5 h post release, likely indicating that cells that were successfully stalled at the beginning of mitosis are yet to undergo cytokinesis. However, 4 h post release there is a significant shift to G_1_ progression, with ∼70% of cell population transitioned through mitosis into the G_1_/G_0_ phase. This result provides the aforementioned time correction between the computational model and measured cyclin protein levels. The rapid change of cell cycle distribution within the first 90 min can be attributed to approximately 20% of cells already in the G_2_/M phase at the point of nocodazole application (data obtained from the untreated control population). This result allows for a more in-depth framing of how cells progress through the cell cycle when translating back to the model. Figure 3 indicates that ∼75% of the cell population is being released into the cell cycle in sync, meaning that when interpreting the subsequent *in vitro* data that the quantised result will be applicable to only 75% of cells.

**Figure 4.**
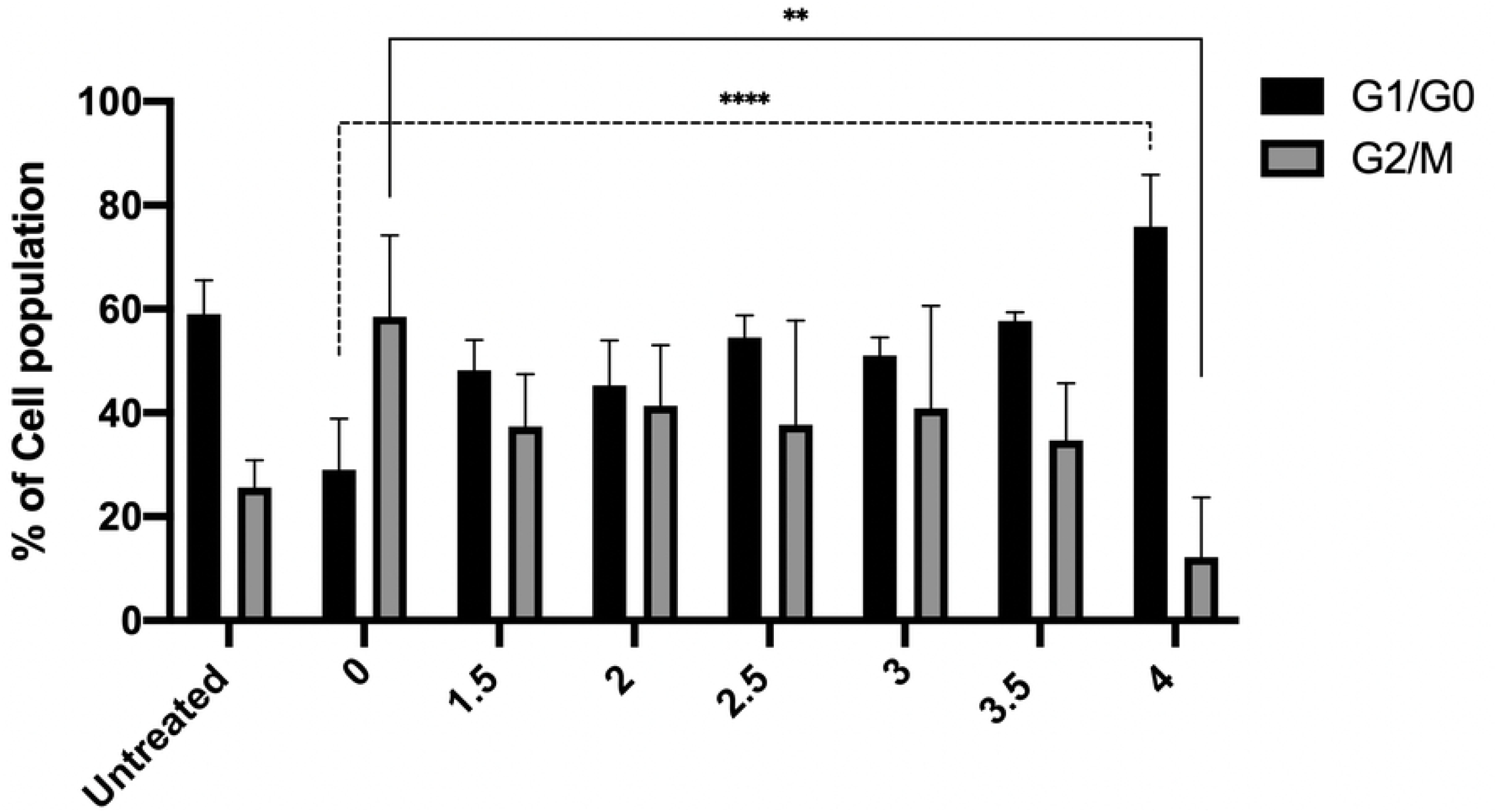
Time dependent release of cells from nocodazole mediated G_2_/M blockage. Typically, by 4 h post release ∼75% of cells have transitioned out of the G_2_/M checkpoint. The increase in G_0/_ G_1_ fraction is highly significant (p<0.0001). As such, this data indicates that a 4 h offset post drug removal should be applied, indicating the beginning of the cell cycle.

The next step was to collect the biological data used for the quantisation of the hybrid model. To fully quantise protein expression levels, standard curves using known concentrations of recombinant cyclin proteins were generated via western blot (Figure 4). From densitometry analysis of the standard curve, equations of best fit were obtained and used in conjunction with the molecular weight of the individual cyclin proteins to calculate absolute concentration of each individual protein.

Whole cell lysates were collected at timepoints spanning a 33 h time period post nocodazole release. This period was selected to ensure complete transition of one cell cycle. Polyacrylamide gels (4-12%) were loaded with 20 μg of whole cell lysate, and probed for each of the aforementioned cyclin proteins and GAPDH as a housekeeping control. Figure 6 details the normalised cyclin protein levels over time, with corresponding representative images each cyclin blot (Figure 6A-D).

**Figure 5.**
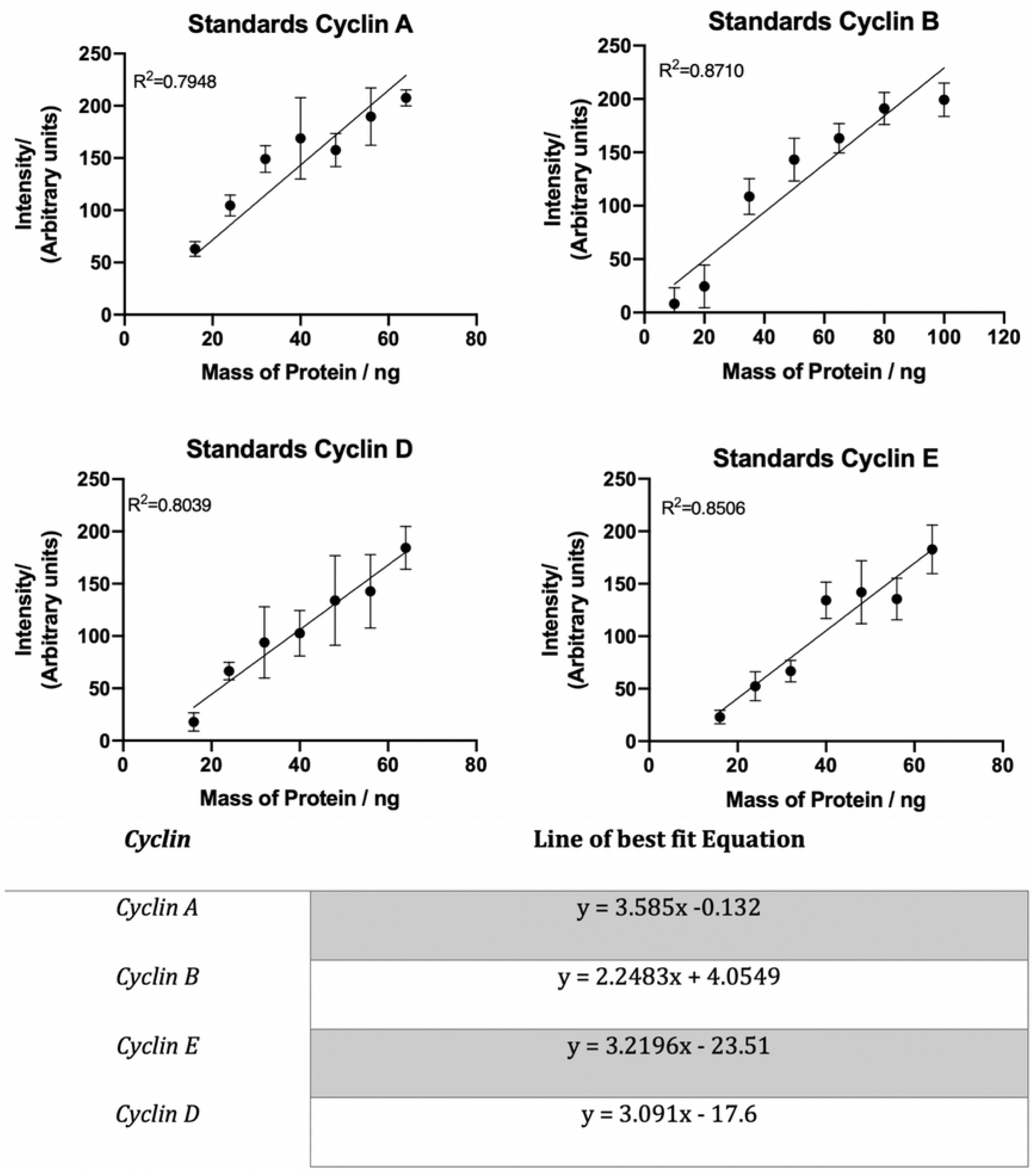
Standard curves of increasing concentrations of recombinant cyclin proteins, used to quantify cyclin expression levels.

**Figure 6.**
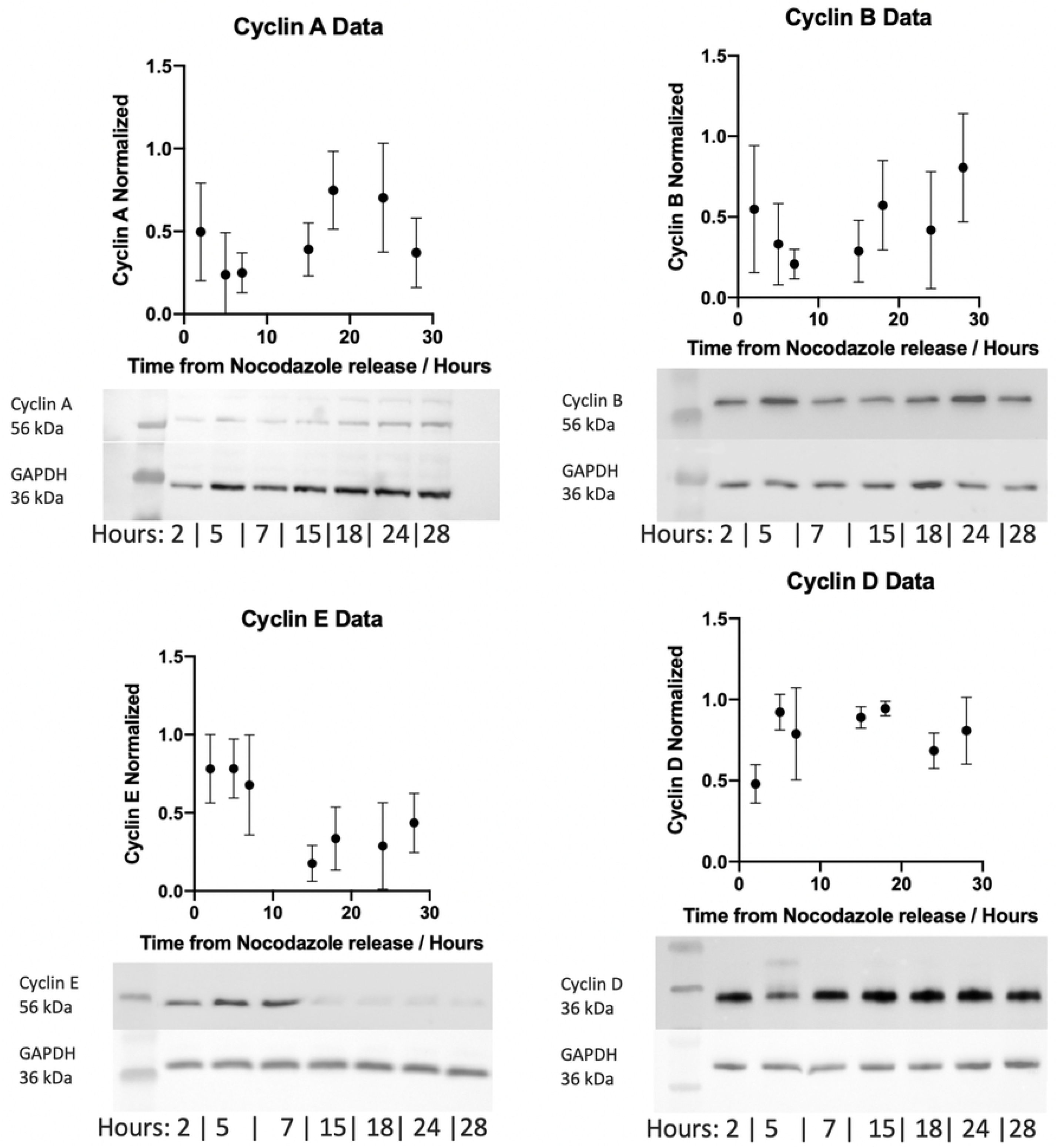
Representative western blots for regulatory cyclin proteins determined by Western blot. The samples were collected over a 28 h period with 40 μg whole cell lysate loaded into each well (n=3).

Western blot band intensities were calculated using the Fiji extension of the ImageJ software. This data was collected using the average pixel colour intensity, with 0 being black and 255 being white. To ensure consistency between each channel the area measured was kept constant. Results where averaged over three independent experiments, with the time series plotted alongside the computational model in Figure 7.

**Figure 7.**
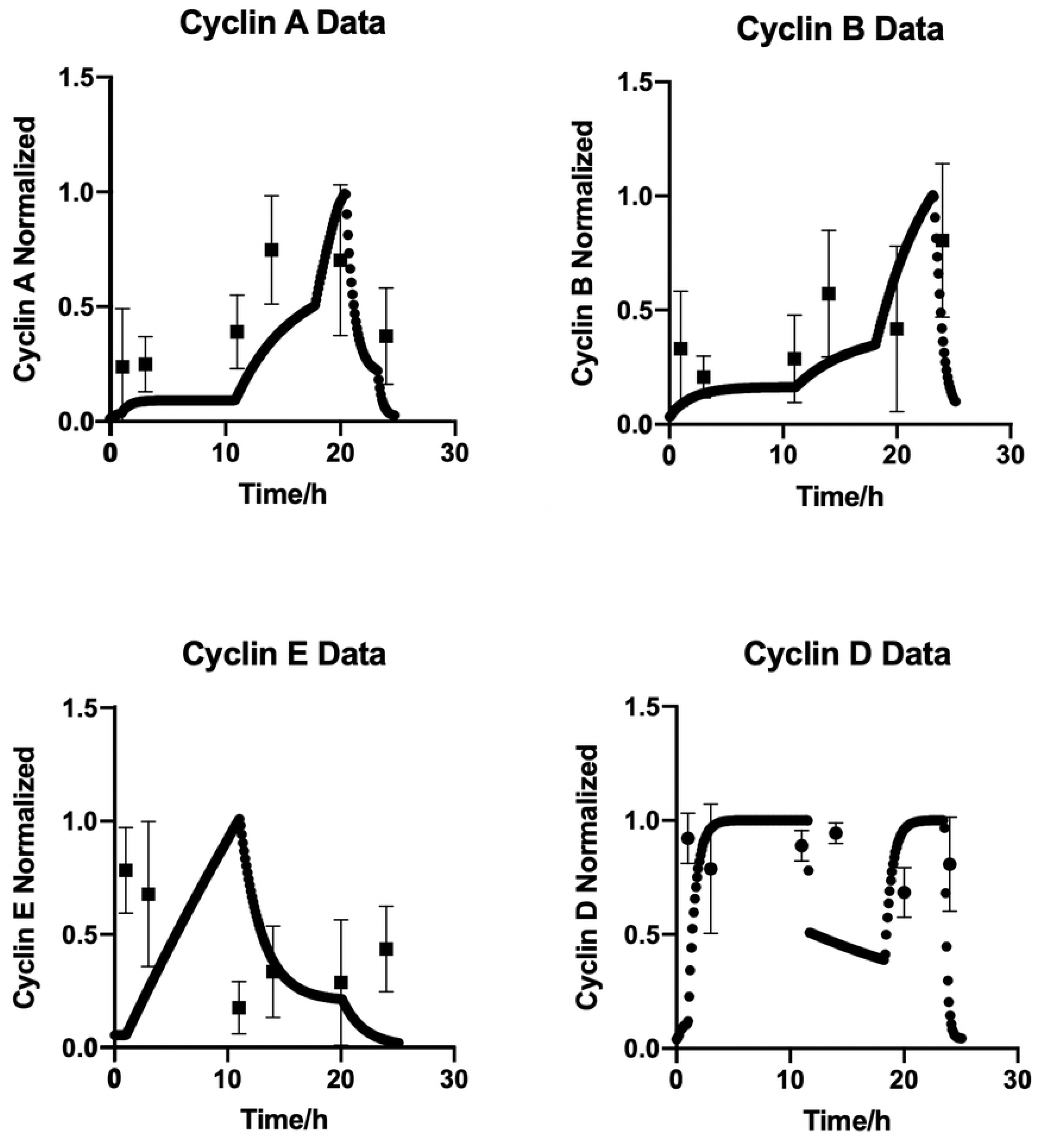
Overlay of the results obtained from both computational modelling and the experimental Western blot data. The biological data in this overlay has been corrected for the 4 h lag observed following nocodazole release. Both datasets have been normalised to allow a more direct comparison.

The computational model used in this study is based on the hybrid model created by Singhania *et al.* (2011).^5^ The model was modified to include a fourth cyclin responsible for cell cycle regulation, cyclin D. This model can be solved in two ways; user defined time intervals, or providing outputs only at the transition between phases. For this study the former was used, providing a more accurate time series to compare against the collected biological data. To calculate the concentration of protein within a single phase three parameters are needed; the initial concentration, the rate of synthesis and the rate of degradation. The initial concentration is that with which the cell enters each phase of the cell cycle with. The rate of synthesis is independent from the protein and is controlled by external factors such as gene expression, while the rate of degradation is dependent on the current level of the protein. This allows for the equations of protein concentrations to take the following form;

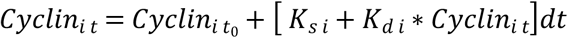

Equation.1 Allows for the concentration of cyclin at a time, *t*, to be calculated. Where *t*_*o*_ is the time at the start of the phase, *i* donates cyclins A,B,E or D, *K*_*s*_ is the rate of protein synthesis and *K*_*d*_ is the rate of protein degradation. The full list of equations can be found in supplementary Table.1 K rates are variables that change as the model progresses thorough the cell cycle. The form taken by K is dependent on the Boolean variables that are either active or inactive during each phase. Their base form are

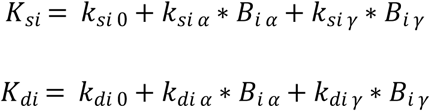

Equation. 2 Where *i* donates the cyclin which *k* is paired with, A B D or E. *K*_*0*_ is the base level of the protein and *k*_*α/γ*_ pair with the Boolean variables that are active in the current cell phase.

As the Boolean values change in every new phase of the cell cycle the *K*_*s*_ and *K*_*d*_ values are re-calculated at the start of each phase. This full Boolean network is shown in the methods section in Table. 2. The simulation produces a clear time series of the levels of cyclin proteins being expressed in a distinct phase of the cell cycle. Figure 7. shows the four time series normalised to their respective highest concentration. This figure also shows the overlay of the time series obtained from cell based experiments shown in Figure 6. Note that as previously stated, the computational model defines time zero as progression out of G_0_ in the cell cycle, while the biological measurements are based on G_2_/M stalling by nocodazole stalling, defining time zero as the start of mitosis. Thus, before direct comparisons can be drawn a correction factor for the time from nocodazole release and to complete mitosis must be applied. This factor was determined as 4 h from the release data presented in Figure 4. The application of this correction and overly of the two data sets can be seen in Figure 7.

**Table. 2.** Boolean variables of key proteins as they change throughout cell cycle phases with corresponding thresholds for each phase.

|  | SCF | CDC25A | CDC20A | CDH1 | CDC20B | CDC25B/C | CDC14 | MYC | Thresholds |
| --- | --- | --- | --- | --- | --- | --- | --- | --- | --- |
| *G0* | 0 | 0 | 0 | 1 | 0 | 0 | 1 | 1 | $C_D > 1.5E^6$ |
| *G1e* | 0 | 1 | 0 | 1 | 0 | 0 | 0 | 1 | $C_E > 4.3E^6$ |
| *G1L* | 0 | 1 | 0 | 0 | 0 | 0 | 0 | 1 | $C_A > 9.35E^5$ |
| *S* | 1 | 1 | 0 | 0 | 0 | 0 | 0 | 0 | $C_E < 1.06E^6$<br>& $t > 5.5$ |
| *G2* | 1 | 1 | 0 | 0 | 0 | 1 | 0 | 1 | $C_B > 6.9E^6$ |
| *Prophase* | 1 | 0 | 0 | 0 | 0 | 1 | 0 | 1 | $t = 0.75$ |
| *Metaphase* | 1 | 0 | 1 | 0 | 0 | 1 | 0 | 0 | $t = 1$ |
| *Anaphase* | 1 | 0 | 1 | 0 | 1 | 1 | 0 | 0 | $t = 0.5$ |
| *Telophase* | 1 | 0 | 1 | 1 | 1 | 0 | 1 | 0 | $C_B < 9.8E^5$ |

The equations from Figure 4 of the standard curves are then applied to the results from Figure 6 to calculate the total amount of cyclin protein in each sample, as a function of absolute cell number, allowing the quantisation of protein molecules per cell throughout specific phases of the cell cycle.

With this definitive information, the computational model can be promoted from calculating a relative level of cyclin proteins to calculating the actual number of protein molecules per cell. The molecular weight of the cyclin proteins are known as cyclin A, B, E and D weigh 52, 48, 50 and 34 kDa respectively.^[24]^ From these weights and the concentrations of the recombinant proteins used to generate the standard curves, the number of protein molecules per cell were calculated varying from 1.2E11(10 ng) to 1.2E^12^ (100 ng). The capabilities of such quantised simulations will allow for the ad hock threshold errors to be more accurately controlled; due to discrete countable systems having an error of 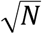, where N represents the number of molecules. The reduction in the errors allows for a stricter control over the model as a whole but also gives insight into the protein levels required to transition through each phase of the cell cycle. With the computational model fully quantised, new simulated data can be calculates and shown in Figure 8.

**Figure. 8.**
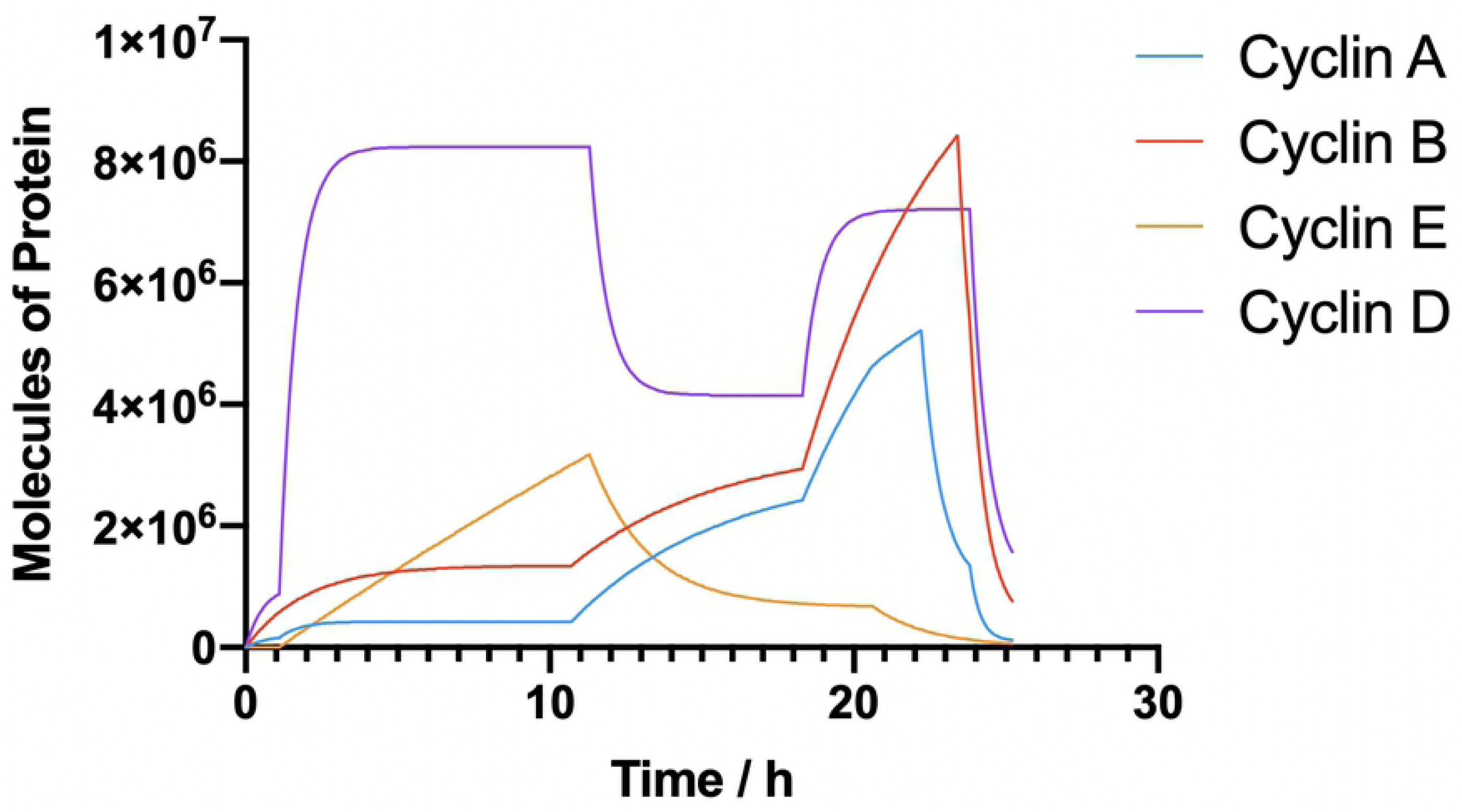
the new output of the code incorporating a quantised number of molecules per cell for all four cyclins.

With the quantised code cyclins A, E and D have a comparatively similar peak values of 5E^6^ molecules per cell with cyclin B expressed highest, yielding approximately 9E^6^ proteins per cell. However, the total number of protein per cell has been calculated to be approximately 2E^9^ meaning that while regulatory proteins are incredibly important to the normal functioning of a cell, they only account for between 0.25 – 0.45% of the total protein content when expressed at their highest concentration.

## Discussion

The importance of a well-informed model cannot be understated. The methodology outlined in this paper allowed for the conversion of a model that worked only within self-constrained concentrations to a model that can be used as a guide to estimate the absolute number of regulatory proteins contained within a single cell, at any one time. To obtain the biological data allowing this conversion, cell cycle synchronisation using nocodazole proved effective, providing fairly uniform release characteristics. Flow cytometry data (Figures 2 & 3) clearly demonstrate the nocodazole induced stalling, plateauing with ∼60% of cells stalled in G_2_/G_m_ at concentrations in excess of 50 μg/ml. Given that HUVEC calls are non-transformed endothelial cells, retaining contact inhibition, we hypothesis, that the remaining cells are likely non-cycling, sitting within G_0._, accounting for the observed G_2_/M plateau.^[25]^ As such, features like contact inhibition highlight the importance of ensuring equivalent confluence between variable *in vitro* cell treatment groups. The absolute values obtained for individual protein molecules during each phase of the cell cycle appear reasonable when compared against total protein concentration for cell lines such as Hela and U20S cells.^[26]^ The total number of protein molecules reported were on the order of 10^9^ molecules per cell. Based on our experimental measurements, the levels of cyclin proteins, even at their most concentrated, is in the order of millions of molecules per cell. Put in context and compared against an abundant protein, such as actin, this measurement is in the region of 2-orders of magnitude lower, with actin reported to have an average concentration of 5E^8^ molecules per cell.^[27]^ This clearly demonstrates that while these regulatory proteins are invaluable to the cell, they convey significant biological effects at comparatively low concentrations.

The model as presented shows good overall agreement between the experimental protein expression data and the computational outputs. This suggests that our hybrid model can provide an accurate output of protein concentration through cell cycle progression. This output also allows for the study of the rates of translation of the proteins. The largest variation in expression was observed with cyclin B, increasing by 4 million molecules over a 5 hour time span. As the cyclin B protein is comprised of 433 amino acids, with a translation rate of ∼10 aa/s, the time approximate time synthesise one cyclin B protein would equate to 43.3 seconds. Therefore, to facilitate the rapid change (5 h) in overall protein expression there must be in the region of 9500 ribosomal sites dedicated to the translation of cyclin B mRNAs.^[28,29]^ The strong agreement across the whole of the cell cycle also indicates the accuracy of the time scale which the model is working across. This ability to have a model that is not computationally intense but provides accurate outputs can, as outlined above, allows for future expansion of the model. With more genes coded into the base Boolean network and more proteins concentrations being calculated, the power of the model could increase, permitting *in silico* analysis of novel drug/radiation treatment combinations.

The main benefit of this model is the fact that it is computationally non-intensive. For future use, this could allow the model to be expanded including processes such as DNA damage and repair or apoptosis, or for it to be absorbed into a larger model without adding excessive computational intensity. Either of these applications would retain the ability to generate protein molecule quantification, while increasing the usefulness of the existing model. In terms of expansion to include cell cycle arrest and DNA damage repair, the gene network that controls the main components of arrest are p53 and p21. These pivotal proteins could be incorporated into to the base Boolean network in Figure 9, thereby expanding the functional relevance of the model .^[30,31]^ With respect to incorporation into an existing model, our hybrid model could be used as a subsection of a more comprehensive tumour vasculature growth model, given that it is based on the proliferation of vascular forming endothelial cells, allowing for variation in the cell cycle of each individual cell rather than being modelled according to set timesteps.^[32]^ The ability to have a computational model that accounts for the study of the full cell cycle while incorporating DNA damage repair and arrest could prove highly valuable in the study of novel cancer therapeutics and the normal tissue response, providing rationalised biological experiments which could limit unnecessary animal use, helping to refine pre-clinical studies in keeping with NC3R guidelines.^[33]^ The primary limitation of this model is that the experimental data used to parametrise the model is based on the proliferation characteristics of normal endothelial cells (HUVEC). Incorporating flexibility to adapt between both tumour and non-tumour would prove highly beneficial, particularly if combined with the ability to model radiation damage for example. This work presented a quantised hybrid model of the cell cycle, which allows for computationally light data collection.

**Figure 9.**
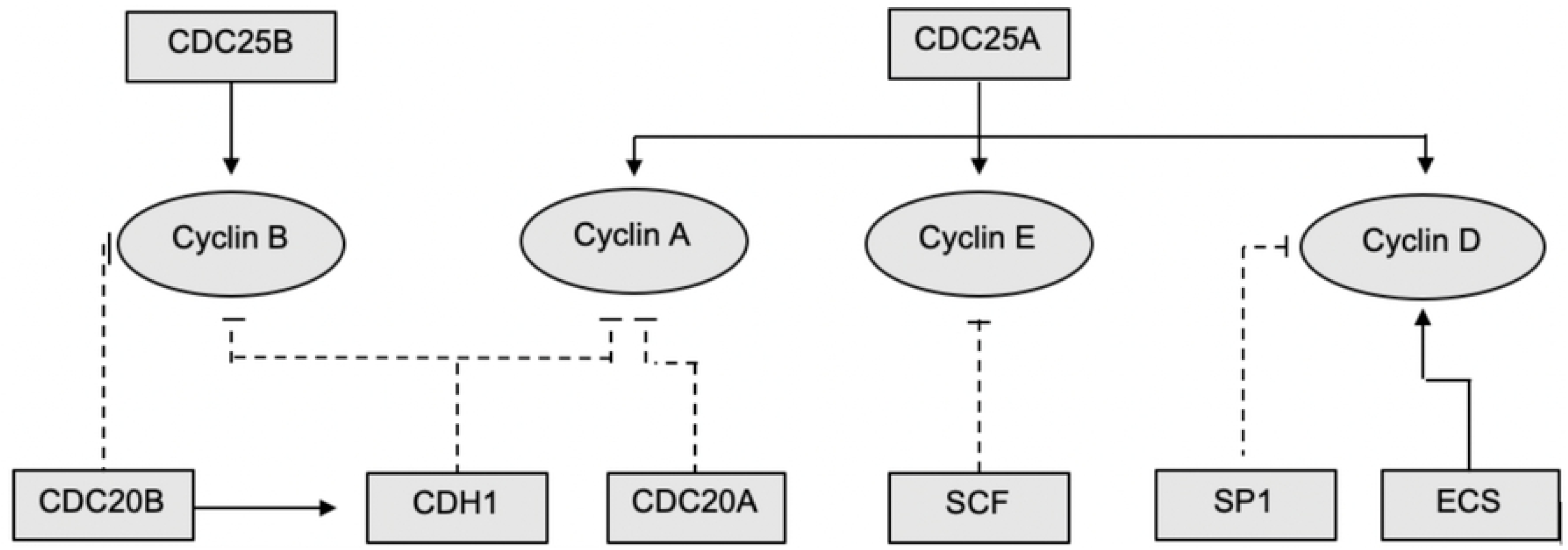
Schematic representation of the interaction network for the promotion and degradation of cyclins and cyclin dependent kinases. Flat ended lines represent suppression and arrows represent up regulation. Boxes denote Boolean variables with circles representing continuous variables within the code.

## Methods

### Computational model

The base of this model is the Boolean network of the main genes responsible for cell cycle regulation. A schematic of this network can be seen in figure 1 with a larger schematic shown below.

This schematic shows the interaction network of the Boolean variables and the alternating regulatory effects which they confer upon the continuous variables represented by cyclins A, B, D and E. The values that the Boolean variables have in each cell phase are given in Table 2. The simulation was solved over 5000 cycles of independent, individual cells. These cells were all initialized with the same concentration of cyclins, cyclin A = B = D = E = 1. However, they were allowed to run through 10 cycles before the 11^th^ cycle was recorded. This provides heterogeneity, as observed in a normal proliferating population, while remaining within the constraints of the model.

At the beginning of each phase new *K*_*s*_ and *K*_*d*_ values are calculated for each cyclin based on Equation 2 and the values from Table 3 as the simulation starts with all values set to 1. The model is set to progress from one phase to the next when the correct threshold conditions are met. For transition from G_1_ to S-phase, the correct thresholds of Cyclin E and Cyclin A must be met. S-phase is both time dependent and cyclin E degradation dependent, while G_2_ and mitosis are governed by cyclin A and cyclin B respectively. In G_1_ the model will solve the time duration required to accumulate sufficient levels of cyclin E. Once this time is calculated, it is used determine the required increase or decrease in expression levels of the remaining three cyclins. Cyclin E concentrations at the end of G_1_ will then become the new initial concentrations for S-phase, with new K rates also determined. This process repeats until the model has solved equations for all the transitions through the cell cycle, reaching the end of the telophase. During telophase, the model will divide the final concentrations of cyclins by two, preparing for cytokinesis, then add a randomised error which will become the new initial condition for the beginning of the next run. After being quantised, the variability in the thresholds where converted from the initial 10% error to be 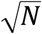 with N being the number of strands of the protein. This 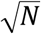 dependence arises from Poisson statistics for counting of discrete entities.

**Table.3.** showing the values for the synthesis and degradation in the mode

| Cyclin A |  | Cyclin B |  | Cyclin E |  | Cyclin D |  |
| --- | --- | --- | --- | --- | --- | --- | --- |
| $K_{sA}$ | $3.85E^5$ | $K_{sB}$ | $9E^5$ | $K_{sE}$ | 5000 | $K_{sD}$ | $1.2E^6$ |
| $K_{sA \text{ cdc}25A}$ | $4.62E^5$ | $K_{sB \text{ cdc}25B/C}$ | $2.1E^6$ | $K_{sE \text{ cdc}25A}$ | $5E^5$ | $K_{sD \text{ cdc}25A}$ | $8.4E^6$ |
| $K_{sA \text{ cdc}25B/C}$ | $1.54E^6$ | $K_{dB}$ | 0.2 | $K_{dE}$ | 0.02 | $K_{dD}$ | 0.802 |
| $K_{dA}$ | 0.2 | $K_{dB \text{ cdc}20B}$ | 1.2 | $K_{dE \text{ SCF}}$ | 0.5 | $K_{dD \text{ myc}}$ | 0.8 |
| $K_{dA \text{ cdh}1}$ | 1.2 | $K_{dB \text{ cdh}1}$ | 0.3 | | | | |
| $K_{dA \text{ cdc}20A}$ | 1.2 | | | | | | |

### Cell culture and Synchronisation

Human umbilical vein endothelial cells (HUVEC) were cultured in ATCC vascular cell basal media supplemented with the ATCC endothelial cell growth kit-BBE. HUVECs were seeded at a density of 3E^5^ in a T25 flask. Cells were seeded 24 h prior to nocodazole (50 μg/ml) stalling at 37°C. After stalling cells were washed twice with sterile PBS (phosphate buffed solution) and then incubated in fresh media for timepoints ranging from 2 h to 33 h. After a predefined period, cells were trypsinised and transferred to a 15 ml flacon tube and centrifuged at 200 g for 5 min. Supernatant was removed and the cell pellet resuspended in 1.2 ml fresh media, and a cell count performed using a Coulter Counter. The remaining cell suspension was again centrifuged, forming a pellet, then stored at -80°C until required.

### Cell lysates

For cell lysate preparation, cell pellets where treated with 100 μL RIPA solution (Thermo Scientific) containing cOmplete™ Protease Inhibitor Cocktail (Sigmal Aldrich). Cell pellets were left on ice for 30 min, vortexed every 10 min, before being returned to -80°C for 10 min. Cell lysate protein levels were then standardised using cell counts via dilution using purified H_2_0. Normalised samples were then combined in equal parts with 2x Laemmli buffer (Sigma Aldrich) prior to denaturing at 95°C for 10 min. Prepared samples were stored at -20°C until required.

### Western Blot

For Western blot analysis, 20 μL of the cell lysates were loaded into each well. Samples were separated by molecular weight via electrophoresis through a 4% – 12% bis-tris gel at 120 mV for 1.5 h. Once separated the proteins where transferred to a nitrocellulose membrane (life sciences) at 35 mV for 2 h. Once transferred, membranes were blocked in 3% weight per volume (w/v) skim milk dissolved in PBS for 1.5 h at room temperature. When blocked, membranes where treated overnight at 4°C with primary antibodies for cyclin A, B, E and D (1:5000). Membranes were then washed three times in 0.1% tween 20 in PBS to remove excess primary antibody before incubation in anti-rabbit secondary antibody (1:5000) for 1 h at room temperature. Antibody chemiluminesence signals were detected using the G-box. Following image acquisition, densitometry was carried out using image-J software.

### Flow Cytometry

Cells where trypsinised at various time points to study the release of cell cycle stalling from nocodazole (90 min, 120 min, 150 min, 180 min, 210 min and 240 min). The cells were washed in PBS after being centrifuged at 250 g for 5 min. Cells were then fixed in 70% ethanol for 2 h at 4°C. Cells were once again washed in PBS before being treated with a propidium iodine (PI) solution (50 μg/mL PI with 1% RNAse) overnight at 4°C. PI florescence was measured over a minimum of 10,000 cells. Samples where gated to remove the debris and DNA doublets, providing cycle separation via total DNA content. The counts of cells in G_0_/G_1_, S-phase and G_2_/M were measured and compared as percentages of the total cell population

### Toxicity Assay

Cells where seeded at a density of 1.2E^4^ cells per well in a 96 well plate, cultured as above and treated with increasing concentrations of Nocodazole (25, 50, 100, 150 and 200 μg/mL) dissolved in media. After 24 h nocodazole treated cells were washed with PBS and assayed for toxicity using AlamarBlue (ThermoFisher). Used at a 1x dilution, cells were incubated in AlamarBlue containing media for 4 h at 37 °C before using florescence measurements as a readout of viability.

## Author Contributions

CE developed the model, carried out all the biological work to validate the model and produced the initial manuscript draft; LB provided wet laboratory and tissue culture support; DG was the secondary project supervisor providing computational modelling support; FC conceived the original idea for the computational model and provided feedback on manuscript drafts; JC was primary supervisor providing support with biological validation, manuscript design and editing.

